# Integrating Orthology and Cross-Species Transcriptomics to Unravel Root Drought Responses in Faba Bean and Maize

**DOI:** 10.64898/2026.09.07.749847

**Authors:** Kübra Arslan, Dagmar van Dusschoten, Agnieszka A. Golicz, Silvia F. Zanini

## Abstract

During soil drying, roots in upper and deeper soil layers experience different hydraulic environments, yet the molecular programs that accompany and potentially drive depth-specific changes in root water uptake capacity remain poorly understood. Here, we address this gap by coupling depth-resolved RNA-seq with previously measured hydraulic uptake profiles in pot-grown faba bean (*Vicia faba*) and maize (*Zea mays*), two crop species with contrasting root architecture, sampled from upper (drying) and lower (comparatively wet) root zones at the onset and after four days of soil drying. Differential expression analysis revealed strongly top-region of the roots dominated transcriptional responses in both species, consistent with the steeper hydraulic challenge in drying upper layers. Maize displayed a TIP- and dehydrin-dominated drought response, whereas faba bean induced NIP-like aquaporins and a broader LEA repertoire, suggesting divergent strategies for root water and cellular stress protection. To compare transcriptome responses across these distally related species, we grouped orthologous genes across 26 species using OrthoFinder. With this approach, we identified 905 drought-responsive orthogroups, of which only 17% were shared between species despite broadly convergent gene onthology (GO) enrichment profiles. Phylogenetic tracing showed that 95.7% of faba bean-specific and 90.3% of maize-specific drought-responsive orthogroups are conserved across monocot and dicot lineages, indicating that species-specific drought transcriptomes arise primarily through differential recruitment of ancestral gene families rather than lineage-specific innovation. These findings define a molecular framework linking root hydraulic architecture to gene regulation under drought and identify conserved transcriptional regulatory hubs as targets for broad-spectrum abiotic stress improvement in both faba bean and maize.

## INTRODUCTION

During drought stress, as soils lose water, soil water potential declines and plants face a cascade of hydraulic challenges: reduced cell turgor (Bartlett et al., 2012), increased risk of xylem dysfunction (Urli et al., 2013), and constraints on phloem transport (Thompson, 2006). The earliest plant response is often stomatal closure, a protective move that also inhibits carbon gain and can force plants toward carbon limitation if drought persists (McDowell, 2011). Yet the severity of water stress experienced by the shoot is not dictated by soil water status alone. It also depends on how the root system’s hydraulic conductance changes during drying, and crucially, where along the soil profile those changes occur.

In drying soil, shallow layers typically dry first while deeper layers remain comparatively wet (Hillel et al., 1976; Kondo et al., 2000). Despite water remaining at depth, plants often still experience substantial drought stress, partly because deeper roots contribute less to whole-plant water uptake under well-watered conditions (Gessler et al., 2022; Passioura, 1983; Prechsl et al., 2015). Additionally, deep roots are commonly less abundant and can be less conductive than shallow roots, making the “deep reservoir” hard to take advantage of when the topsoil becomes hostile (Müllers et al., 2022). A drought-tolerant plant is therefore not merely characterized by deep rooting depth, but by its capacity to dynamically reallocate the hydraulic contribution of distinct root zones over time.

Recent non-invasive profiling of root water uptake during early soil drying revealed that total root system conductance can decline steeply even under moderate drought, and that this decline is not spatially uniform (Van Dusschoten et al., 2020). Conductance in shallow, drying layers tends to decrease, while in some cases conductance in deeper, wetter layers can increase, partially compensating for losses near the surface (Müllers et al., 2023). These patterns imply active, localized regulation: reducing uptake in the driest zones may help preserve hydraulic continuity at the soil-root interface, while enhancing intrinsic conductivity at depth can improve access to remaining water (Müllers et al., 2023). Candidate mechanisms include changes in aquaporin activity (Johnson et al., 2014; McLean et al., 2011; Rodríguez-Gamir et al., 2019) and deposition of apoplastic barriers such as suberin (Barrios-Masias et al., 2015). However, the molecular programs that accompany and potentially drive this depth-specific hydraulic redistribution remain poorly resolved, especially in a comparative framework across species with contrasting root architectures and water uptake strategies.

Here, we address this gap by linking depth-resolved transcriptomic responses to previously observed depth-dependent hydraulic shifts during early soil drying (Müllers et al., 2023). We sampled roots from the upper (drier) and lower (wetter) soil layers after sustained drying (Day 4) or under control conditions in two crop species with contrasting root system organization: faba bean (taproot-dominated) and maize (fibrous). We generated RNA-seq profiles for each depth and condition, performed differential expression analyses, and used an orthology-based strategy to enable cross-species comparison at the level of shared gene families and conserved functional modules. This design allowed disentangling (i) responses to soil drying, (ii) spatial responses associated with local soil water status, and (iii) species-specific versus conserved regulatory programs.

We hypothesize that upper roots exposed to soil drying will show transcriptional signatures consistent with reduced radial water flow where as lower roots in wetter layers are expected to show signatures consistent with maintaining or enhancing water uptake capacity. By comparing these transcriptomic reprogramming between maize and faba bean through orthogroup-level analyses, we aim to identify conserved drought-response “cassettes” that operate across architectures, as well as species-specific strategies that may contribute to elucidating differences in the ability to sustain deep-water uptake during early drought.

## MATERIALS AND METHODS

### Experimental Setup and Growth Conditions

Twelve plants of each species were grown in 50cm long tubes with an inner diameter of 81mm and a soil depth of 45cm. Six plants of each species were not subjected to drought and used as controls. The soil mix, consisting of 80% agricultural loamy sand was mixed with 20% coarse sand (see Müllers, 2022 for details). At sowing the pots were watered up to 25% (580 mL) soil water content (SWC) with 0.3% HakaPhos Red nutrient solution. After three weeks another 200 mL of the same nutrient solution was applied. In between, the plants were regularly watered making sure the SWC never dropped below 15%, a level at which full transpiration is maintained. The potted plants were grown in a climate chamber with 50% humidity at a constant 20°C. Furthermore, plants were grown in 14h daylight and 10h nighttime. During daytime light was modulated in 2h blocks between high light and low light, between 1000 and 150 µmol.m^-2^s^-1^ respectively. The light modulation was used to quantify zonal, relative root water uptake (van Dusschoten 2020).

Plants were grown for 5 weeks in well wetted conditions. Four days prior the planned harvest half of the plants received their last water such that the initial water content was 20% SWC. The control plants were watered up to 25% and received additional water one day before harvest.

### Soil water content measurements

The six plants of each species selected for drought stress were placed in the Soil Water Profiler (SWaP) during a four day period. The SWaP measures SWC profiles of the 45cm columns in 1cm steps using a moveable sensor that resides on the outside of the tube of the potted plants (for a detailed description, see van Dusschoten et al. 2020). The instrument also measures the local water changes with very high precision. By using light modulation the rapid plant responses can be separated from the slower through-soil water mobility such that the local water concentration changes can be converted to plant driven root water uptake (RWU). This information, combined with the measured SWC content was used here to define the drought level of the zones that were excavated. Similarly the total change of the SWC, Utot or total RWU was used to identify the stress level of the plants. The SWCs of plants not subjected to drought were determined gravimetrically.

As the SWaP instrument can only measure four plants at a time the complete experiment was split into three blocks each with two faba bean and two maize plants that were droughted and two of each species used for control. At the time of each of the first two harvests new plants were sown, limiting the total time of this experiment to 18 weeks.

### RNA-seq Sampling, Sequencing and Data Processing

At the end of the four-day drought period, roots at a depth of 5-10cm (top) and 25-30cm (bottom) were collected in liquid N_2_ for subsequent RNA.seq. Total RNA was extracted with Zymo *Quick*-RNA Plant Kit (Zymo Research) and sent for pair-ended Illumina library preparation and sequencing to Novogene(Novogene Gmbl). Raw sequencing reads were subjected to quality assessment using FastQC v0.12.1 (*Babraham Bioinformatics - FastQC A Quality Control Tool for High Throughput Sequence Data*, n.d.) to evaluate overall sequencing integrity. Adapter sequences and low-quality bases were trimmed using Trimmomatic v0.39 (Bolger et al., 2014), employing default parameters optimized for paired-end RNA-seq data.

### Transcript Assembly, and Quantification

High-quality reads were aligned to the reference genomes of faba bean (Jayakodi et al., 2023) and maize (Hufford et al., 2021) using HISAT2 v2.2.1 (Kim et al., 2019) with strand-specific settings (--rna-strandness RF) and an intron length threshold (--max-intronlen 10000) appropriate for plant genomes. Aligned reads were sorted and indexed using SAMtools (Li et al., 2009) sort and flagstat to facilitate downstream processing. Transcript assembly was performed with StringTie v2.2.3 (M. Pertea et al., 2015), using conservative parameters (--rf -j 3 --conservative -m 500) to minimize fragmentation. A consensus transcriptome was generated using the --merge option (-c 100 -m 100) to integrate individual assemblies. Non-annotated transcripts were systematically renamed according to species designation (Vfaba for *Vicia faba* and Zmays for *Zea mays*), and only the longest isoform per gene was retained using custom scripts. Transcript sequences were extracted from the reference genome using GffRead (G. Pertea & Pertea, 2020) with -w option, and coding sequences were predicted with TransDecoder (*DataLad Repository: Datasets.Datalad.Org/Shub/Sghignone/TransDecoder*, n.d.), ensuring accurate annotation of potential open reading frames. To refine transcript annotations, reference genome annotations were intersected with the newly assembled transcripts using BEDTools (Quinlan & Hall, 2010) with intersect -wa option, allowing the identification of novel transcripts absent from the reference annotation. These unique transcripts were integrated with existing genome annotations, and the final curated annotation was used for transcript re-quantification via StringTie (M. Pertea et al., 2015) (-eB). Gene- and transcript-level count matrices were generated using the prepDE utility from StringTie.

### Differential Expression Analysis

Gene expression matrices derived from the re-quantification step were filtered to exclude low-expression transcripts. Genes were retained if they (i) exhibited a minimum expression threshold of five normalized counts and (ii) were detected in at least two biological replicates per condition and region. Differential expression analysis was conducted using DESeq2 (Love et al., 2014), employing a generalized linear model (GLM) with the design formula ~region + condition to account for both regional and drought stress effects. In addition, independent DE analyses were performed within each root region to specifically assess local transcriptomic responses to drought.

### Functional Annotation, Gene Ontology (GO) Enrichment and Phylogenetic Analyses

Using officially annotated and curated transcripts from the transcript assembly and quantification step, Blast2GO (Conesa et al., 2005) and InterproScan (Jones et al., 2014) were run to get functional annotation of the genes using the databases of *Oryza sativa* and *Arabidopsis thaliana* (default parameters were used in OmicsBox). Using filtered gene count tables from differentially expression analysis step as background genes, differentially expressed genes examined for enrichment by using topGO (Alexa & Rahnenfuhrer, 2023). The semantic similarity between enriched GO terms were measured using GOSemSim (Yu, 2020) package with semantic similarity threshold of 0.6. For selected water stress-related gene families, candidate proteins were identified by combining domain-based annotation with BLAST-based similarity searches against *A. thaliana* proteins (Altschul et al., 1990; Camacho et al., 2009). Aquaporin candidates were identified using the Pfam aquaporin domain PF00230 and the corresponding InterPro domain IPR000425, and the resulting candidates were merged with BLASTp hits against *A. thaliana* aquaporin proteins (Quigley et al., 2002). LEA candidates were identified using a custom database composed of LEA1–LEA6, seed maturation protein (SMP), and dehydrin sequences were retrieved(Hundertmark & Hincha, 2008).

BLASTp hits were filtered using a minimum sequence identity of 30% and a minimum alignment coverage of 80%. Candidate proteins were assigned gene symbols according to the best matching *A. thaliana* reference genes and *A. thaliana* nomenclature. For each gene family, protein sequences were aligned using MUSCLE5 (Edgar, 2022), and phylogenetic dendrograms were reconstructed with the maximum-likelyhood (for aquaporins) and neighbor-joining (for LEAs) method implemented in MEGA software (Kumar et al., 2018).

### GO Enrichment and Orthogroup-Based Mapping (GO×OG)

Orthologous genes between faba bean and maize were identified using OrthoFinder v 3.0.1b1 (Emms & Kelly, 2019). To improve orthology inference, protein sequences of 23 phylogenetically intermediate and closely related species of faba and maize were used (Supp. Table 1) after checking BUSCO duplication scores (Supp. Table 1) to prevent unnecessary duplication leading to false positive copy number variation inference. Genes associated with significantly enriched BP terms (p < 0.05) were selected as candidates. These genes were then mapped across species based on their orthogroup assignments to identify orthologous relationships. For each gene within these (GO×OG), the corresponding molecular function (MF) annotation was also retrieved, enabling cross-species comparison of both biological processes and associated molecular functions using custom build scripts.

## RESULTS

### SWaP guided-sampling and drought stress measurements

At the time of harvest total RWU had decreased on average to 0.3 times RWU at the start of the drought period for both species (Fig. 1). For faba bean the upper layer contained 5.6 ± 0.1 % of water (−0.16 MPa) and the lower layer 6.6 ± 1% (−0.11 MPa) (Fig. 1A and B). For maize the upper layer contained 4.5 ± 0.8 % of water (−0.27 MPa) and the lower layer 5.8 ± 0.5% (−0.15 MPa) (Fig. 1C and D). Both species had an identical lower RWU at the upper layer that was reduced to 26%, whereas for the lower layer faba bean RWU was reduced to 43% and for maize to 78% of the initial values.

**Figure 1.**
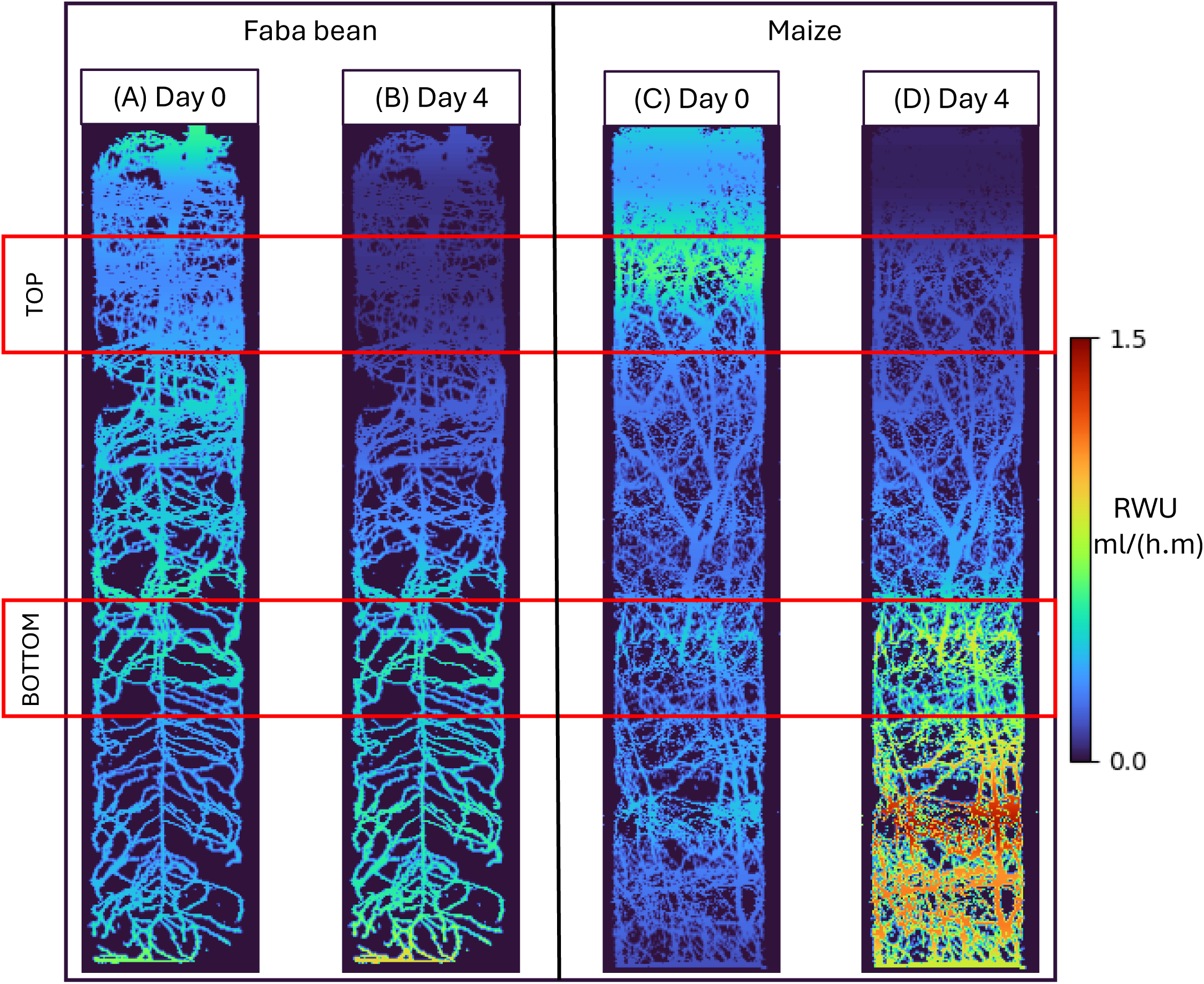
Exemplary Maximum Intensity Projections of MRI images of roots of faba bean (A and B) and maize (C and D) at DAS 37 (Days After Sewing). The roots are color coded with Root Water Uptake (RWU) per root length (ml/(h.m)) amplitudes acquired with the Soil Water Profiler (SWaP) on DAS 39 (A and C) and DAS 43 (B and D). Red boxes are indicating the depth of excavation. Root lengths were determined via NMRooting (van Dusschoten et al, 2016) on the MRI images. At DAS 39 the soil was relative wet (between 17 and 25 % soil water content (SWC) top to bottom respectively) which allows for unhampered root water uptake. At DAS 43 SWC had declined to about 7% at the top and 15% at the bottom resulting in compensatory RWU patterns especially so for the Maize plants. Here, no separate MRI images could be acquired after the imposed drought as roots needed to be harvested. Potential root growth is therefore not corrected for in these images but was found to be limited in an earlier study (Müllers, 2023).

### Sequencing, Assembly and Annotation

A total of 48 RNA-seq libraries (PE 150 bp) were generated across faba bean and maize samples representing top and bottom root regions under control and drought conditions. For faba bean, the average number of sequenced reads per sample were around 44.3 million, with a GC content of 41–43%. For maize, sequencing yielded on average 45.25 million reads per sample, with GC content ranging from 51% to 54% (Supplementary Table 1-2). High-quality reads alignment efficiency was consistent across all samples, with faba bean libraries achieving an average total mapping rate of 92.7% ± 0.5%, and maize libraries reaching an average of 86.4% ± 0.7% (Supp. Table 3). The proportion of uniquely (primary) mapped reads was similarly high, averaging 91.5% for faba bean and 84.7% for maize, ensuring accurate transcript assembly and quantification (Supp. Table 3).

Transcript assembly followed by comparison with the corresponding reference genome annotations identified a total of 1,912 novel transcripts in maize (2.57% of the final merged transcriptome) and 1,321 novel transcripts in faba bean (3.42%). The final merged transcriptomes comprised 74,451 transcripts in maize and 38,573 transcripts in faba bean, combining both annotated and novel transcripts.

### Depth-Resolved Transcriptional Responses to Drought

Differential expression analysis was performed to assess the independent effects of root region (top or bottom) and drought treatment across all samples. Additionally, separate differential expression analyses were applied within each root region to identify region-specific transcriptional responses to drought stress. After read-count filtering, 39,388 genes for maize and 33,645 genes for faba bean were kept for downstream analyses.

#### Faba Bean

Drought induced a substantial transcriptional response in faba bean roots. Across both root regions combined (6 controls vs 6 under stress condition), 1,030 genes (3.06% of total gene number) were differentially expressed (|log_2_FC| > 2, padj < 0.05; Fig. 2), indicating a broad activation of stress-responses.

**Figure 2.**
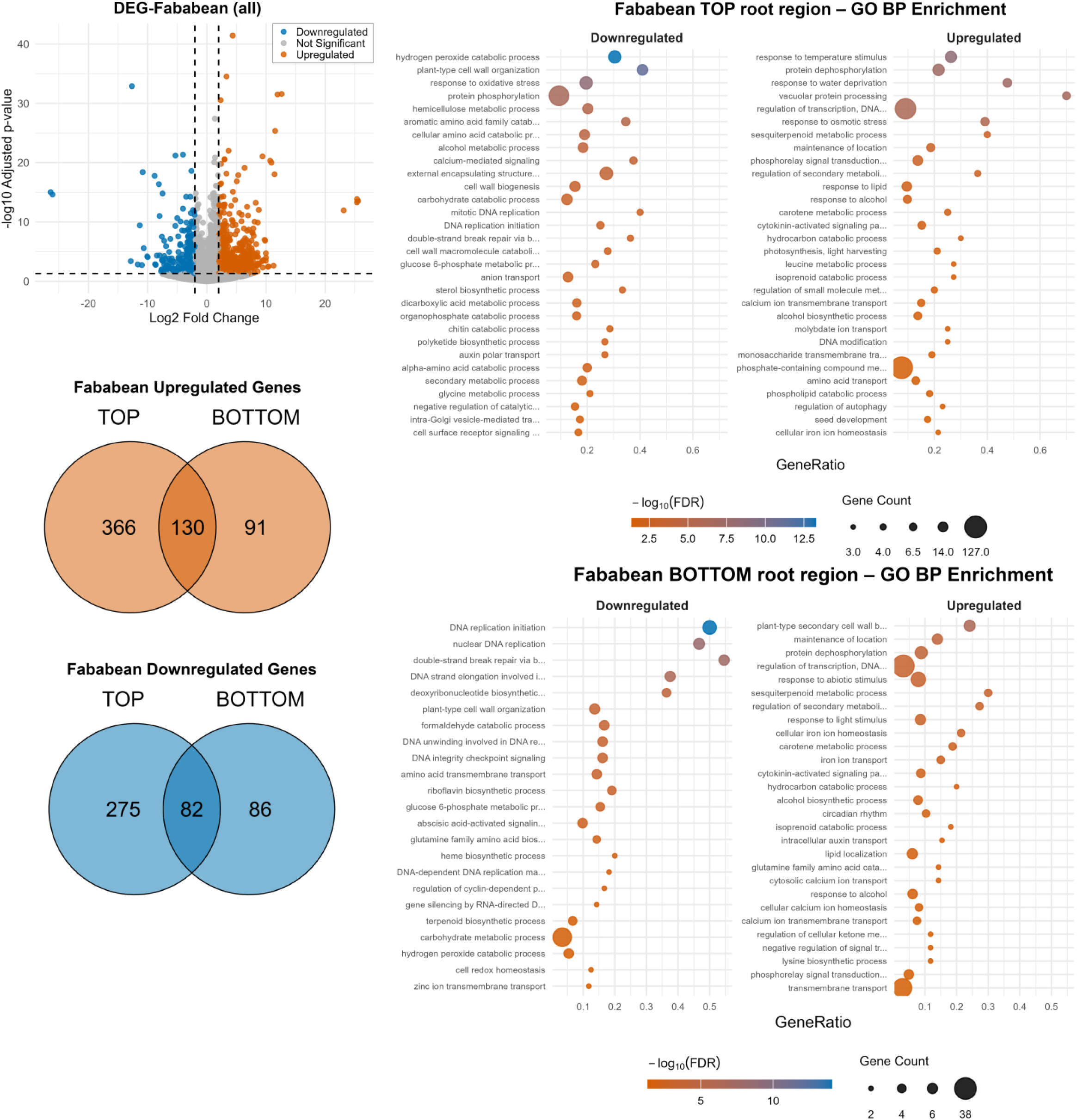
Depth-resolved transcriptional response to drought stress in faba bean roots. (Top left) Volcano plot of differentially expressed genes (DEGs) across both root regions combined (|log_2_FC| > 2, padj < 0.05); orange: upregulated, blue: downregulated, grey: not significant. (Middle left) Venn diagram of upregulated DEGs between top and bottom root regions. (Bottom left) Venn diagram of downregulated DEGs. (Top right) GO biological process enrichment in the top root region, shown separately for downregulated (left) and upregulated (right) gene sets; bubble size represents gene count, colour indicates −log₁₀(FDR). (Bottom right) GO biological process enrichment in the bottom root region.

The spatial decomposition of this response revealed a pronounced top-dominance. In the upper root region, exposed to the strongest drying, 496 genes were upregulated and 357 were downregulated compared to control conditions (Fig. 2). The bottom region showed an attenuated response, with only 221 upregulated and 168 downregulated genes. Approximately half of the differentially expressed genes detected in the deeper (wetter) zone were not shared with the shallow zone (91 upregulated, 86 downregulated), confirming distinct transcriptional responses between the two sections.

GO biological process enrichment in the top root region reflected signatures of acute water stress (Fig. 2). Among upregulated processes we detected an enrichment for response to water deprivation, response to osmotic stress, and regulation of transcription pointed to ABA-mediated drought signaling. Notably, phosphate-containing compound metabolism was the GO term enriched with the largest gene count (127 genes). Among downregulated processes in the top region, protein phosphorylation and plant-type cell wall organization featured prominently, consistent with a reduction in growth-associated signaling and wall remodeling activity under stress (Fig. 2). While regulation of transcription remained a major upregulated category, the bottom root region additionally enriched for response to abiotic stimulus, transmembrane transport, and calcium ion transmembrane transport, suggesting the activation of ion transport and signaling machinery even in the comparatively wetter lower zone. A notable feature of the deeper root regions downregulation was a strong enrichment of DNA replication initiation and nuclear DNA replication processes, suggesting a suppression of cell proliferation programs in the lower root despite its relatively favorable water status, consistent with a systemic growth.

#### Maize

Drought also elicited a broad transcriptional response in maize roots, with 1,034 genes (2.62% of total gene number) differentially expressed across both regions (|log_2_FC| > 2, padj < 0.05; Fig. 3).

**Figure 3.**
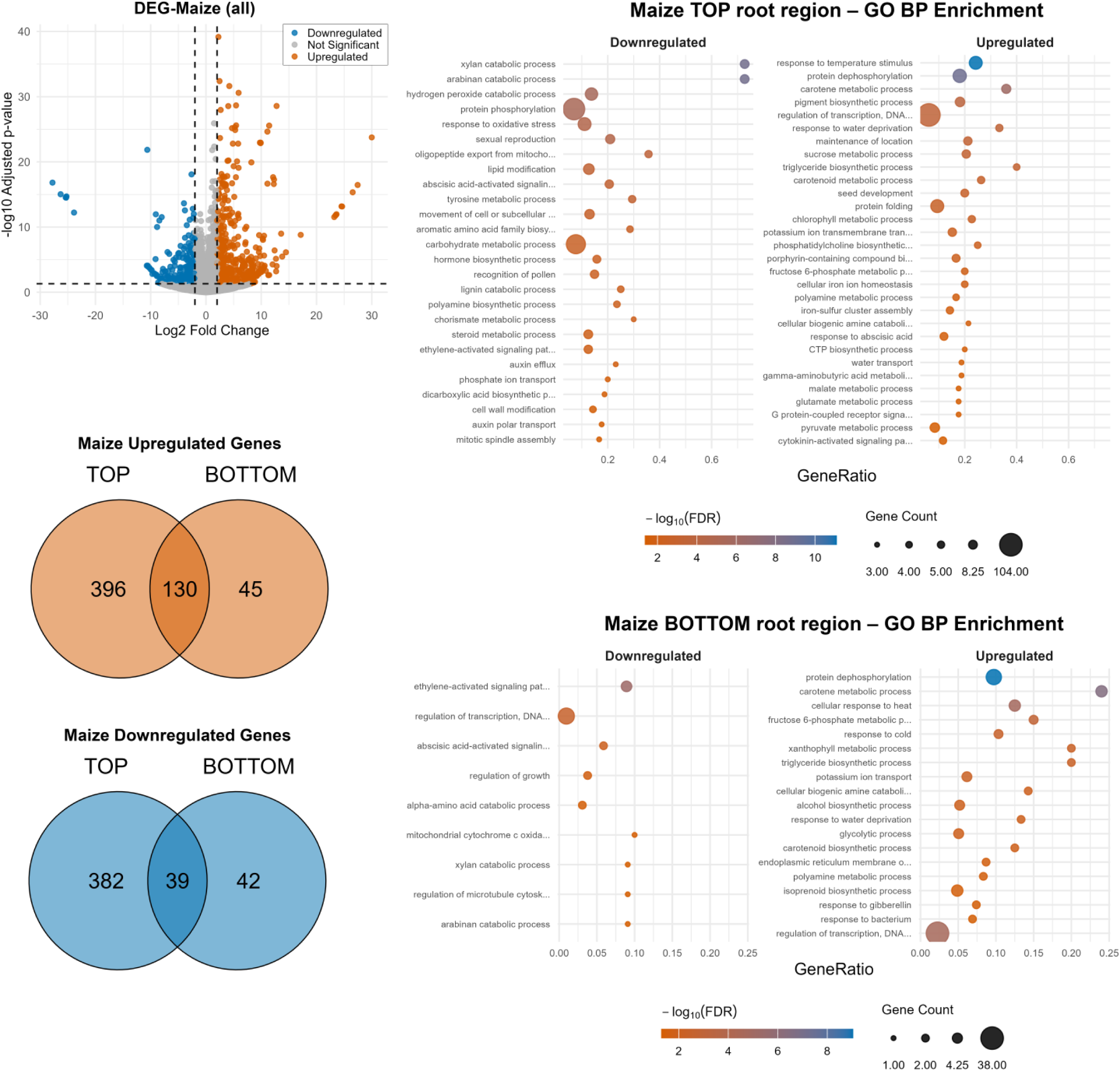
Depth-resolved transcriptional response to drought stress in maize roots. (Top left) Volcano plot of DEGs across both root regions combined (|log_2_FC| > 2, padj < 0.05); orange: upregulated, blue: downregulated, grey: not significant. (Middle left) Venn diagram of upregulated DEGs. (Bottom left) Venn diagram of downregulated DEGs. (Top right) GO biological process enrichment in the top root region; bubble size represents gene count, colour indicates −log₁₀(FDR). (Bottom right) GO biological process enrichment in the bottom root region.

Compared to faba bean, the maize response was even more top-dominated, with the top region exhibiting 526 upregulated (396 top-specific plus 130 shared) and 421 downregulated genes (Fig. 3). The bottom region showed a reduced reprogramming, with only 175 upregulated and 81 downregulated genes.

GO enrichment in the maize top region revealed a mixed signature of stress activation and metabolic reorganization (Fig. 3). Upregulated processes included response to water deprivation, response to temperature stimulus, and regulation of transcription, matching patterns observed in faba bean. Additionally, an enrichment for water transport, protein folding, sucrose metabolic process, and potassium ion transmembrane transport related terms pointed to an active management of cellular osmolyte balance and protein homeostasis. Triglyceride and phosphatidylcholine biosynthetic processes terms were also enriched among upregulated genes, suggesting membrane lipid remodeling as part of the drought response in the maize upper root.

Among downregulated processes, protein phosphorylation and carbohydrate metabolic processes controlled, a pattern shared with faba bean, alongside xylan and arabinan catabolic processes, suggesting a reduced hemicellulose turnover consistent with dampened cell wall dynamics under stress (Jardine et al., 2022; Le Gall et al., 2015). Abscisic acid-activated signaling terms appeared enriched among downregulated genes in the top region, potentially reflecting a post-signaling attenuation of pathway activity rather than absence of ABA-mediated response (Ng et al., 2014; Yang et al., 2016).

The maize bottom region upregulated genes were found enriched in carotene and carotenoid metabolic processes, xanthophyll metabolic process, and fructose 6-phosphate and glycolytic processes, pointing toward increased carbon mobilization and antioxidant carotenoid production in the wetter lower zone. Response to water deprivation was also enriched among upregulated genes in the bottom region, suggesting that even the comparatively less-stressed lower roots perceive and respond to the systemic drought signal. Downregulation in the deeper roots was limited to a small number of processes, including ethylene-activated signaling and regulation of growth, consistent with reduced ethylene-mediated growth promotion as part of a systemic drought response (Růžička et al., 2007).

### Gene Family-Level Transcriptional Responses to Drought

To complement the genome-wide differential expression analysis, we focused on selected gene families representing major components of drought adaptation, including Aquaporins, as key membrane water transporters, and LEA/dehydrin, associated with dehydration-protection. To complement the initial functional annotation we additionally performed custom search and phylogenetic classification and integrated the results with expression profiles (Fig. 4-5 and Supp. Fig. 1-2).

**Figure 4.**
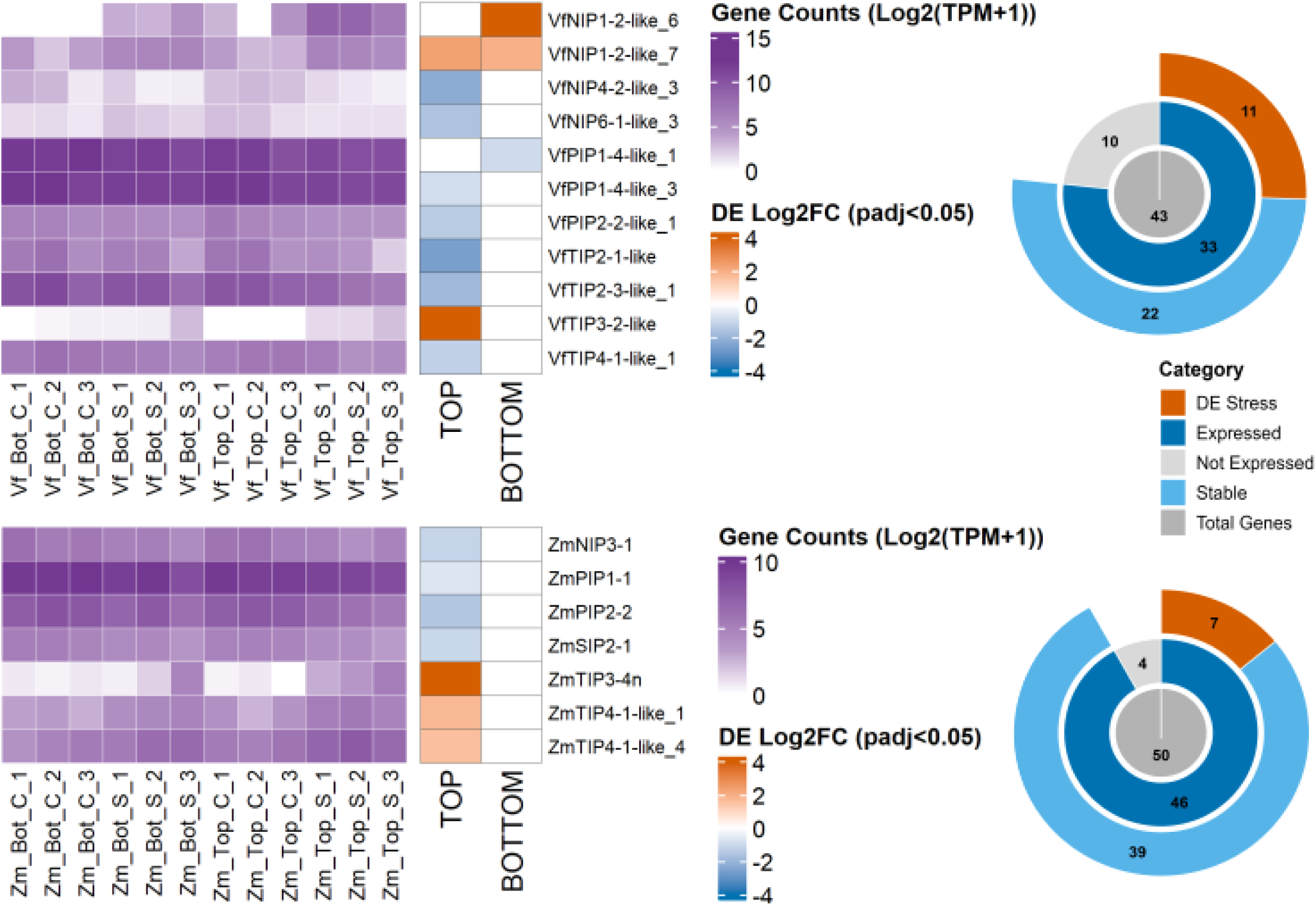
Aquaporin gene expression and drought-responsive aquaporin subclasses in faba bean and maize roots. Heatmaps show log_2_(TPM + 1) expression values of selected aquaporin genes across control and drought samples from bottom and top root regions in faba bean and maize. Adjacent heatmaps show region-specific differential expression as log_2_ fold change under drought in top and bottom root regions; only significant changes are shown for padj < 0.05. Donut plots summarize the number of total aquaporin genes annotated, expressed genes, non-expressed genes, drought-responsive genes, and stably expressed genes in each species.

**Figure 5.**
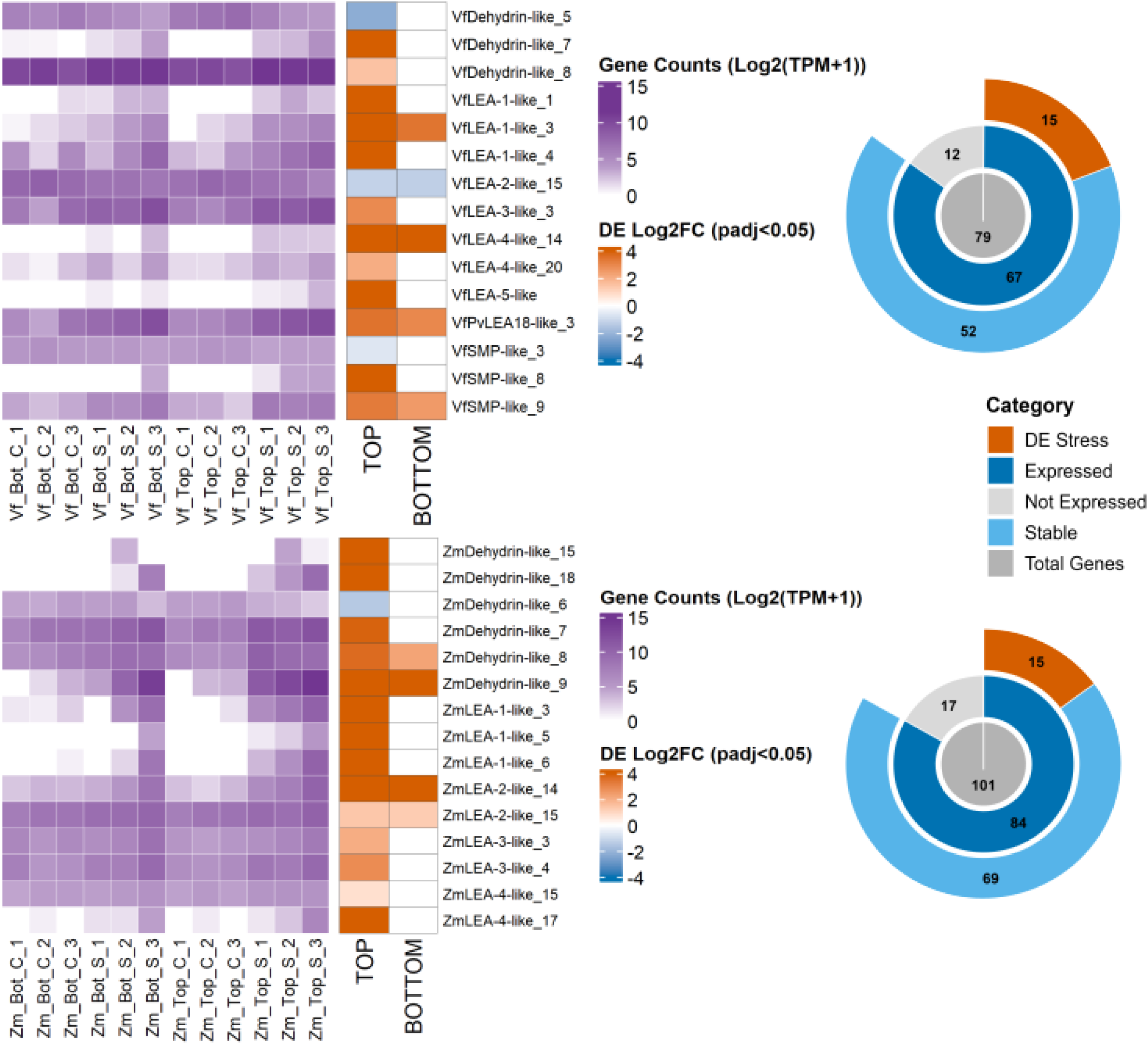
LEA-related gene expression and region-specific drought responses in faba bean and maize roots. Heatmaps show log_2_(TPM + 1) expression values of selected LEA-related genes across control and drought-stressed samples from bottom and top root regions in faba bean and maize. Adjacent heatmaps show region-specific differential expression as log_2_ fold change under drought in top and bottom root regions; only significant changes are shown for padj < 0.05. Donut plots summarize the number of total LEA-related genes annotated, expressed genes, non-expressed genes, drought-responsive genes, and stably expressed genes in each species.

### Aquaporins

A total of 50 genes were annotated as aquaporins in maize, out of which 46 (92%) were found expressed in our samples. Seven maize aquaporins (14%) were found differentially expressed during drought stress compared to control conditions (Fig. 4, Supp. Fig. 1 and Supp. Table 5).The TIP (Tonoplast Intrinsic Protein) type aquaporins were the only upregulated ones and two PIPs (Plasma membrane Intrinsic Protein), a SIP (Small basic Intrinsic Protein) and a NIP (Nodulin 26-like Intrinsic Protein) type aquaporins were downregulated at the top region of the root, there were no differentially expressed gene found at the bottom region of the root (Fig. 4). In faba bean, 43 aquaporin genes were annotated by both BLASTP and Blast2GO domain search (Fig. 4, Supp. Fig. 1 and Supp. Table 4). Only 33 of them (~77%) were expressed and 11 of them (~26%) were found differentially expressed. In contrast to maize, faba bean exhibit differential stress response at the bottom region of the roots. Two NIP1 were significantly upregulated and one PIP type aquaporin were downregulated. At the top of the roots, similar to maize, a TIP3 and NIP1 were upregulated and three TIP, two PIP and 2 NIP types of aquaporins were downregulated (Fig. 4).

### LEAs

Late embryogenesis abundant (LEA) proteins and their subfamily member dehydrins showed high expression in both root regions of maize and faba bean with generally stronger expression in the top region across conditions (Fig. 5, Supp. Fig. 2 and Supp. Table 7). In total, 79 and 101 LEA family proteins were annotated for faba bean and maize respectively. Out of which 15 proteins differentially expressed under stress conditions for both plants. In contrast to maize, faba bean had significant expression of SMPs in both regions or the roots. Maize did not show any differential regulation of SMPs and had LEA1-4 and dehydrin families upregulated in both regions of the roots (Fig. 5, Supp. Fig. 2 and Supp. Table 6).

### GO Enrichment and Orthology Mapping

To place the species-specific gene family responses identified above within a broader evolutionary framework and assess whether the observed transcriptional divergence reflects lineage-specific gene content or differential recruitment of shared ancestral genes, we performed a genome-wide orthology analysis across 26 plant species. Orthologous gene grouping across 26 species (maize, fababean and 24 intermediate species) using OrthoFinder v2.5 yielded 32,319 orthogroups encompassing 811,457 of 896,217 genes (90.5%), with a mean orthogroup size of 25.1 genes. Species-specific orthogroups accounted for only 9,842 groups (5.1% of all assigned genes), reflecting the broad conservation of gene families across clades.

Of 905 orthogroups containing drought-responsive genes, 392 (43%) were exclusive to faba bean, 359 (40%) to maize, and only 154 (17%) were shared between both species (Fig. 6B-C),. Top enriched GO terms, including regulation of transcription, phosphorylation, and response to oxidative stress, were represented across all three orthogroup categories, yet were driven by different gene sets depending on the species (Fig. 6A), suggesting convergent pathway-level responses underpinned by divergent gene sets. Regulation score analysis (UP-DOWN defined as the number of upregulated minus downregulated genes per orthogroup) revealed a subset of orthogroups with strong concordant upregulation in both species, notably those associated with transcriptional regulation and protein dephosphorylation, while orthogroups linked to oxidative stress and hydrogen peroxide catabolism showed strong downregulation specific to *V. faba* (Fig. 7A, Supp. Fig. 3). Tracing the phylogenetic distribution of species-specific orthogroups across 26 species showed that 95.7% of V. faba-specific and 90.3% of Z. mays-specific orthogroups were conserved across both monocot and dicot lineages (Fig. 6B). Only 9.7% of maize-specific orthogroups were monocot-restricted, with dicot-restricted orthogroups being rare in *V. faba*. These findings indicate that the observed species-specific drought transcriptional reprogramming in maize and faba bean arise primarily through differential recruitment of ancestral gene families rather than lineage-specific gene innovation, suggestiong regulatory divergence as the main driver of species-specific stress adaptation.

**Figure 6.**
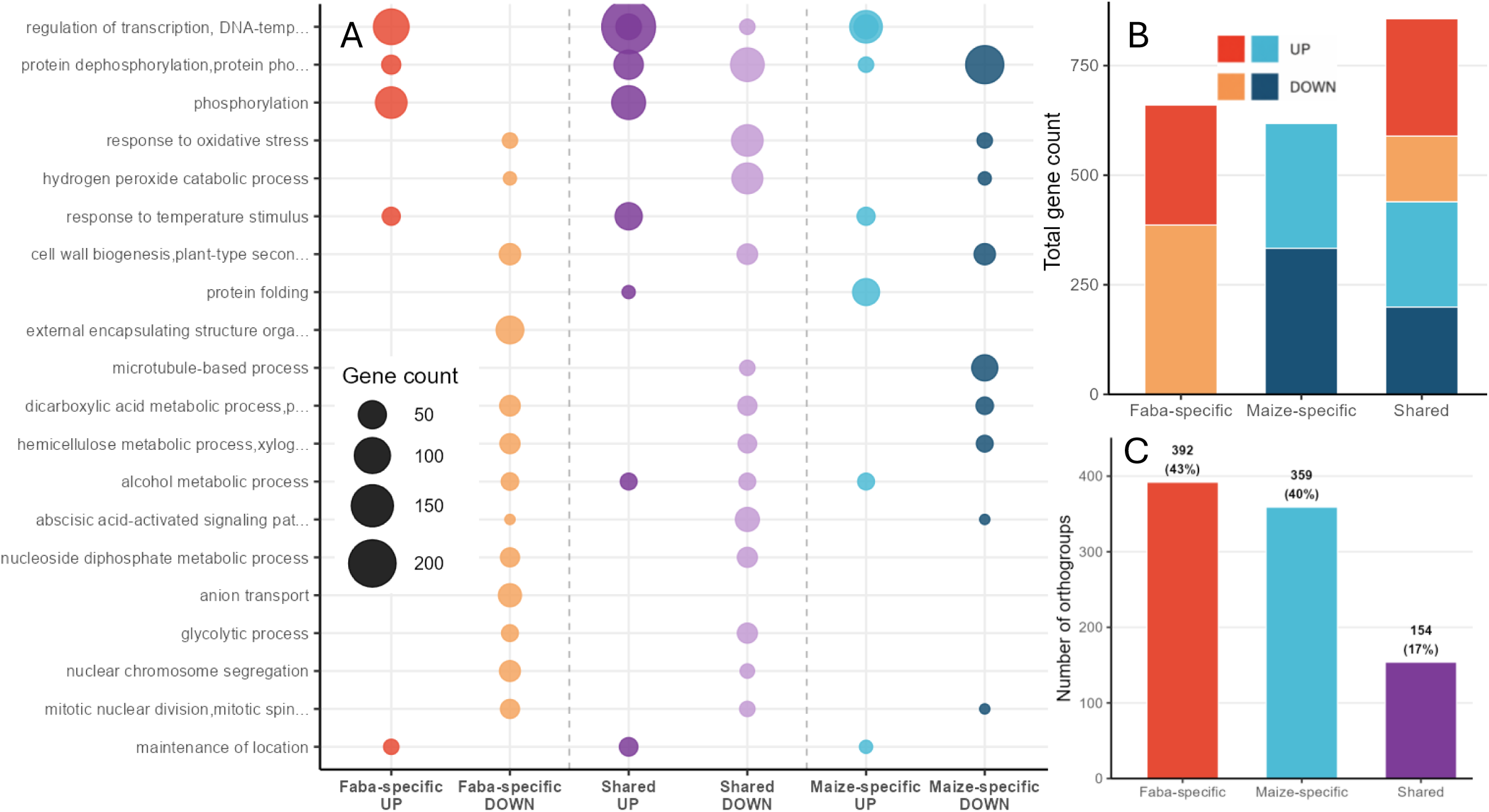
Orthogroup-level comparative analysis of drought-responsive genes across faba bean and maize. (A) Bubble plot of top GO biological process terms enriched among drought-responsive orthogroups, stratified by species specificity and regulation direction (Faba-specific UP/DOWN, Shared UP/DOWN, Maize-specific UP/DOWN); bubble size represents gene count. (B) Stacked bar chart of total gene counts in drought-responsive orthogroups across faba-specific, maize-specific, and shared categories, coloured by species and regulation direction. (C) Bar chart of faba-specific, maize-specific, and shared drought-responsive orthogroup counts; values above bars indicate counts and percentages.

**Figure 7.**
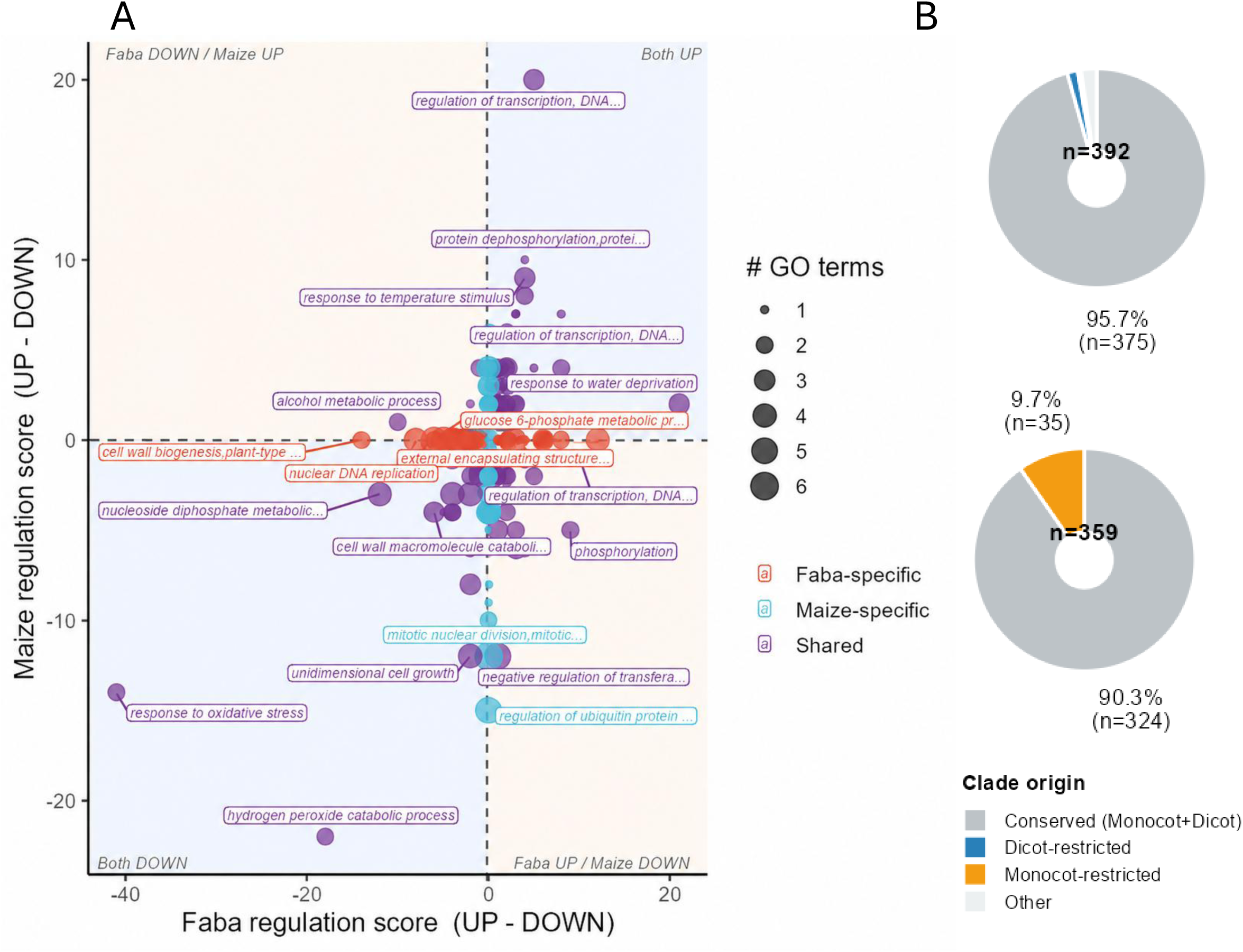
Regulation directionality and phylogenetic conservation of drought-responsive orthogroups. (A) Regulation score scatter plot comparing UP − DOWN gene balance per orthogroup in faba bean (x-axis) vs. maize (y-axis). Each bubble represents one orthogroup; size reflects number of associated GO terms; border colour indicates species assignment (faba-specific: red/orange; maize-specific: cyan; shared: purple). Upper-right quadrant: concordant upregulation in both species; lower-left: concordant downregulation; key orthogroups are labelled. (B) Phylogenetic conservation of faba bean-specific (upper donut) and maize-specific (lower donut) drought-responsive orthogroups. Colours indicate clade origin: conserved across monocots and dicots, monocot-restricted, dicot-restricted (blue, trace in faba bean), and other (light grey).

## DISCUSSION

In this study, physiological measurements were used to guide root sampling at different soil depths, enabling a cross-species comparison of transcriptional responses associated with different physiological responses to drought. The spatial distribution of transcriptional responses reported here is consistent with the depth-dependent hydraulic phenotype documented in the same experimental system by Müllers et al. (2023). In both species, differential expression was strongly top-dominated, with the drying upper root region producing several-fold more DE genes than the wetter bottom region (Fig 1 and 2). A similar depth-dependent shift was observed in field-grown maize, where water deficit reduced root biomass in the upper 0–30 cm but increased root biomass in the 30–100 cm soil layer, indicating greater root investment in deeper soil regions under drought (Heinemann et al., 2026). This spatial difference mirrors the 66–72% reduction in Kroot measured in upper soil layers versus the stable or transiently increasing conductance in lower layers (Müllers et al., 2023), and reflects the steeper water potential gradient experienced by shallow roots. The top-specific enrichment of response to water deprivation, osmotic stress, and ABA-mediated signaling in both species thus aligns with the degree of hydraulic challenge at each depth rather than representing a uniform whole-root response. This interpretation is consistent with the view that root systems are functionally heterogeneous, with different root classes and soil depths contributing differently to water uptake under drought (Comas et al., 2013).

A hypothesis from the hydraulic data was that reduced upper root conductance is partly driven by aquaporin architecture and increased suberization, while the transient conductance increase in deep maize roots could reflect locally enhanced aquaporin activity (Comas et al., 2013; Lindh et al., 2014; Müllers et al., 2023). Our gene family expression analysis not only confirms the role of aquaporins in mediating upper-root conductance reduction, but also identifies a set of novel gene family-level differences between species and root regions that collectively shows the contrasting deep-root strategies of faba bean and maize.

In maize, only specific TIP-type aquaporins were found upregulated. TIPs mainly mediate water transport across the vacuolar membrane and contribute to cytosolic osmotic balance, while some members can also transport small solutes such as urea, ammonia, hydrogen peroxide, and glycerol. Their induction may therefore reflect broader regulation of intracellular water and solute balance under drought (Zou et al., 2024). In faba bean, the drought response showed divergent aquaporin pattern, with upregulation of two NIP-like proteins rather than the TIP-dominated response observed in maize. NIPs have been shown to facilitate the transport of water as well as small neutral solutes, with substrate specificity depending on their pore selectivity residues and subgroup classification (Zhang et al., 2022). Functional characterization of soybean nodulin 26, the protein after which the NIP aquaporin subfamily was named, showed that it transports both water and glycerol and likely contributes to osmotic regulation rather than uncontrolled water loss (Dean et al., 1999). Therefore, the induction of NIP-like genes in faba bean may reflect drought-induced remodeling of membrane permeability and water/solute balance. In contrast, the maize TIP-based response, may indicate a more directly water-storage-related adjustment through the tonoplast.

Beyond aquaporin regulation, LEA gene families revealed a strong region-specific drought response in both species with differences in the LEA subclasses involved in maize and faba bean (Fig. 4). In maize, several dehydrin-like, LEA1-like and LEA2-like genes were found significantly upregulated during drought predominantly in the top root region, indicating activation of a dehydration-protection program in the tissue most exposed to water scarcity. In contrast, the bottom region showed only a limited LEA response, suggesting that severe water scarcity was mainly restricted to the upper root region and had not yet reached the deeper roots, although these tissues may also rely on different protective mechanisms. Faba bean displayed a similarly top-biased LEA response, but with a broader set of LEA subclasses, including LEA1-, LEA4-, LEA5-, LEA18-like and SMP-like genes.Notably, SMP-like genes were drought-responsive only in V. faba, suggesting that faba bean may recruit seed maturation-associated desiccation-protection components in vegetative root tissues under drought (Battaglia & Covarrubias, 2013; Hundertmark & Hincha, 2008). This pattern indicates that both species activate LEA-mediated cellular protection under drought, but differ in the composition of the responding LEA subclasses.

To determine whether the species-specific transcriptional responses reflected different gene content or different regulation of shared gene families, we compared drought-responsive genes at the orthogroup level. The low overlap of drought-responsive orthogroups between maize and faba bean agrees with other comparative transcriptomic studies showing that species exposed to similar stress can share some responses, while most expression changes remain species-specific (Dias et al., 2022; Garassino et al., 2024; Hartmann et al., 2022; Lee et al., 2025; Wang et al., 2023). In two legume species, *Parkia platycephala* and *Stryphnodendron pulcherrimum* grown on iron-rich canga substrate, Dias et al. found a small set of conserved ortholog responses, with most expression patterns differing between species and distinct biological processes affected in each species (Dias et al., 2022). In our data, the shared enriched GO terms were not a result of the same drought-responsive genes being differentially expressed in both species, but rather that maize and faba bean appear to affect conserved functional categories through different gene sets (Fig. 5A-C).

A recent cross-species drought network analyses in Japonica rice, indica rice, wheat, maize, and sorghum,, where drought resistance was described as a network-level trait with both conserved and species-specific components (Deng et al., 2026). In that study, conserved modules included ABA-related PP2C regulation and redox-related gene families, while other regulatory programs differed between crop lineages (Deng et al., 2026). Our results fit this pattern, with transcription factor and protein dephosphorylation orthogroups showing the clearest shared upregulation between maize and faba bean, while oxidative stress- and hydrogen peroxide-related orthogroups showed shared downregulation, indicating that both conserved activation and suppression are part of the drought response (Fig. 6A). In contrast, orthogroups associated with external encapsulating structure organization were specifically downregulated in faba bean, indicating a species-specific suppression of cell wall-related processes under drought. A similar reduction in cell wall-related transcription has been reported in banana, where most drought-responsive genes associated with cell wall metabolism, including cellulose synthases, expansins and pectin-modifying enzymes, were downregulated under water deficit (Muthusamy et al., 2016).

Interestingly, species-specific drought-responsive orthogroups were mostly present across both monocots and dicots, indicating that differences observed between maize and faba bean are not primarily due to lineage or species specific genes or gene families. Instead, both species reprogram conserved gene families under drought conditions. Similar patterns have been reported in drought-related legume genomic studies, where conserved gene content was associated with species-specific changes in stress-related pathways (Chakraborty et al., 2023; Dias et al., 2022; Moghaddam et al., 2021). Overall, these results suggest that maize and faba bean differ mainly in how conserved gene families are regulated during drought, rather than in the presence or absence of drought-related gene families.

These findings carry practical implications for drought improvement in both crops. The dominance of regulatory divergence over gene-content divergence indicates that the molecular raw material for enhanced drought adaptation, the gene families themselves, is largely shared and phylogenetically ancient. Variation in cis-regulatory elements and trans-acting factors that determines which family members are recruited under which conditions is therefore likely to be determining the divergence in responses observed in this study and to be proming targets for tolerance improvement. For faba bean, a crop that lags substantially behind maize in genomic and breeding infrastructure but now has a chromosome-scale reference genome (Jayakodi et al., 2023), the orthogroup framework established here provides a route to leveraging knowledge from better-studied systems. The identification of conserved transcriptional regulatory hubs, points to candidate nodes where intervention may yield broad benefits across drought, salinity, and heat stress, which frequently co-occur under field conditions (Bokszczanin et al., 2013; Jiang et al., 2025).

## CONCLUSION

We analyzed depth-resolved transcriptional responses to soil drying in the root systems of *V. faba* and *Z. mays* and integrated them with previously measured depth-dependent hydraulic phenotypes in the same experimental system. Transcriptional responses were strongly top-dominated in both species, consistent with the greater hydraulic challenge in drying upper soil layers, and included gene family-level signatures, aquaporin downregulation, peroxidase-driven apoplastic barrier reinforcement. The absence of a clear deep-root aquaporin transcriptional response in maize, despite hydraulically documented conductivity increases, points to post-translational regulation as the likely mechanism for rapid conductance adjustment in wetter soil layers. At the orthogroup level, only 17% of drought-responsive gene groups were shared between species, with 90–96% of species-specific orthogroups conserved across monocot and dicot lineages, indicating that differential recruitment of ancestral gene families, rather than lineage-specific gene innovation, is the primary driver of species-specific drought adaptation. Together, these findings define a molecular framework linking root hydraulic architecture to gene regulation under drought, with direct relevance for identifying targets in drought-resilience breeding of both faba bean and maize.

## Supplementary Fıgure Legends

Supp. Fig. 1. Phylogenetic tree of Aquaporins in A. thaliana, faba bean and maize

Supp. Fig. 2. Phylogenetic tree of LEAs in A. thaliana, faba bean and maize

Supp. Fig. 3. GO molecular function enrichment among drought-responsive orthogroups in V. faba-specific, shared, and Z. mays-specific categories. Bubble size represents gene count.

## Supplementary Table Legends

Supplementary Table 1. FastQC quality metrics for V. faba RNA-seq libraries. Per-file output for all 12 samples; columns show sample name, % duplicates, GC content, average read length, and total sequences (M Seqs). Samples sequenced across multiple lanes are listed separately. Abbreviations: Bot = bottom root region; Top = top root region; C = control; S = drought-stressed; 1–3 = biological replicate.

Supplementary Table 2. FastQC quality metrics for Z. mays RNA-seq libraries. Same format as Supplementary Table 1. Abbreviations as above.

Supplementary Table 3. HISAT2 alignment statistics for all V. faba and Z. mays samples. Columns show sample name, total input reads, total mapping rate (%), and primary (unique) mapping rate (%). Multi-lane samples were merged prior to alignment. Abbreviations: Bot = bottom; Top = top; C = control; S = stressed.

Supplementary Table 4. Aquaporin gene annotations and expression in faba bean.

Supplementary Table 5. Aquaporin gene annotations and expression in maize.

Supplementary Table 6. LEA protein gene annotations and expression in faba bean.

Supplementary Table 7. LEA protein gene annotations and expression in maize.

Supplementary Table 8. Plant species included in the OrthoFinder v2.5 comparative genomic analysis. Listed with phylogenetic clade assignment (Monocot, Dicot, or Basal Angiosperm), BUSCO protein completeness score against the embryophyta_odb10 benchmark (n = 425; C: complete [S: single-copy, D: duplicated], F: fragmented, M: missing), and reference.

