## Supplementary figures and images for "Integrating Orthology and Cross-Species Transcriptomics to Unravel Root Drought Responses in Faba Bean and Maize"

### Supplementary Figures

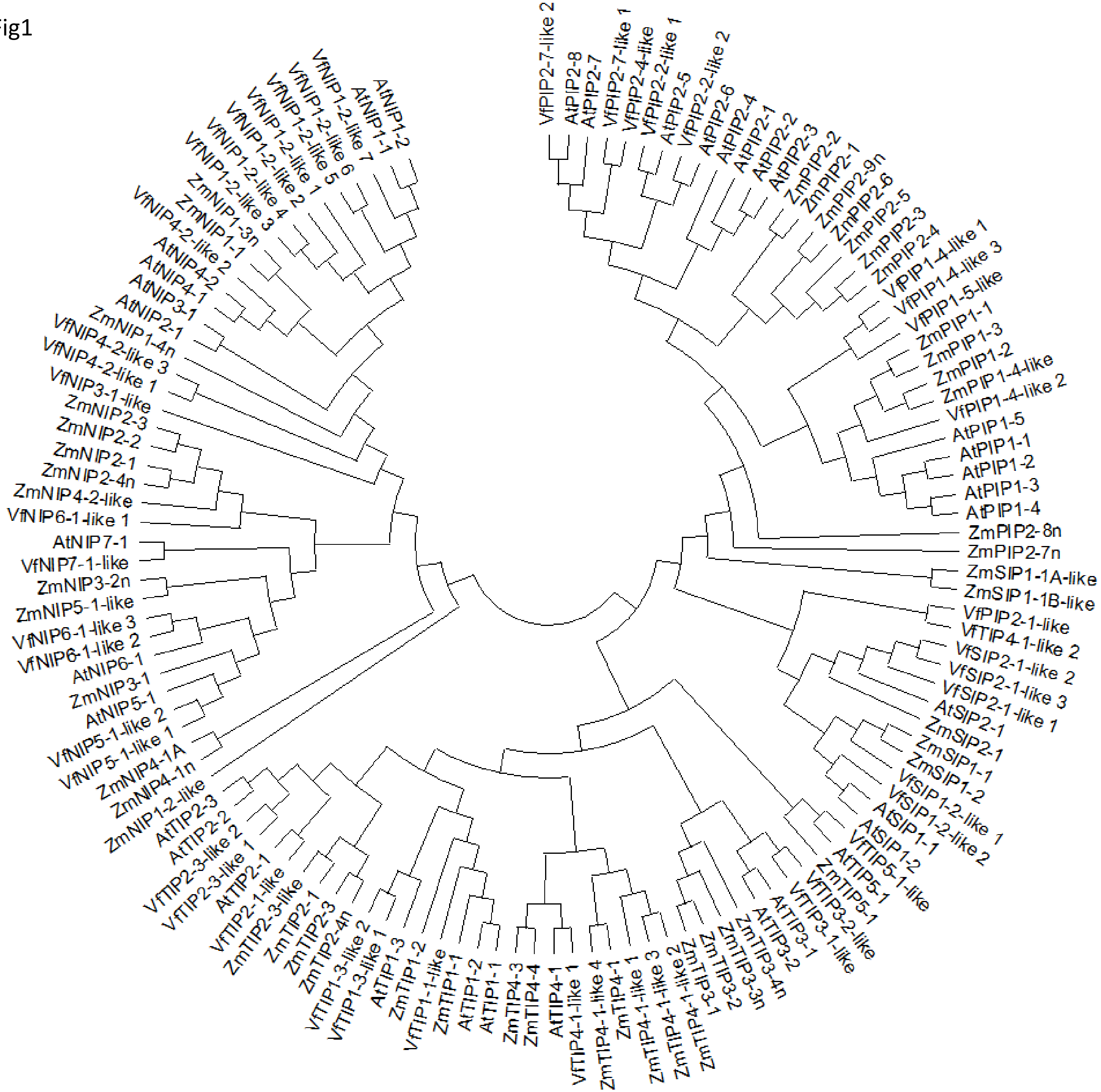

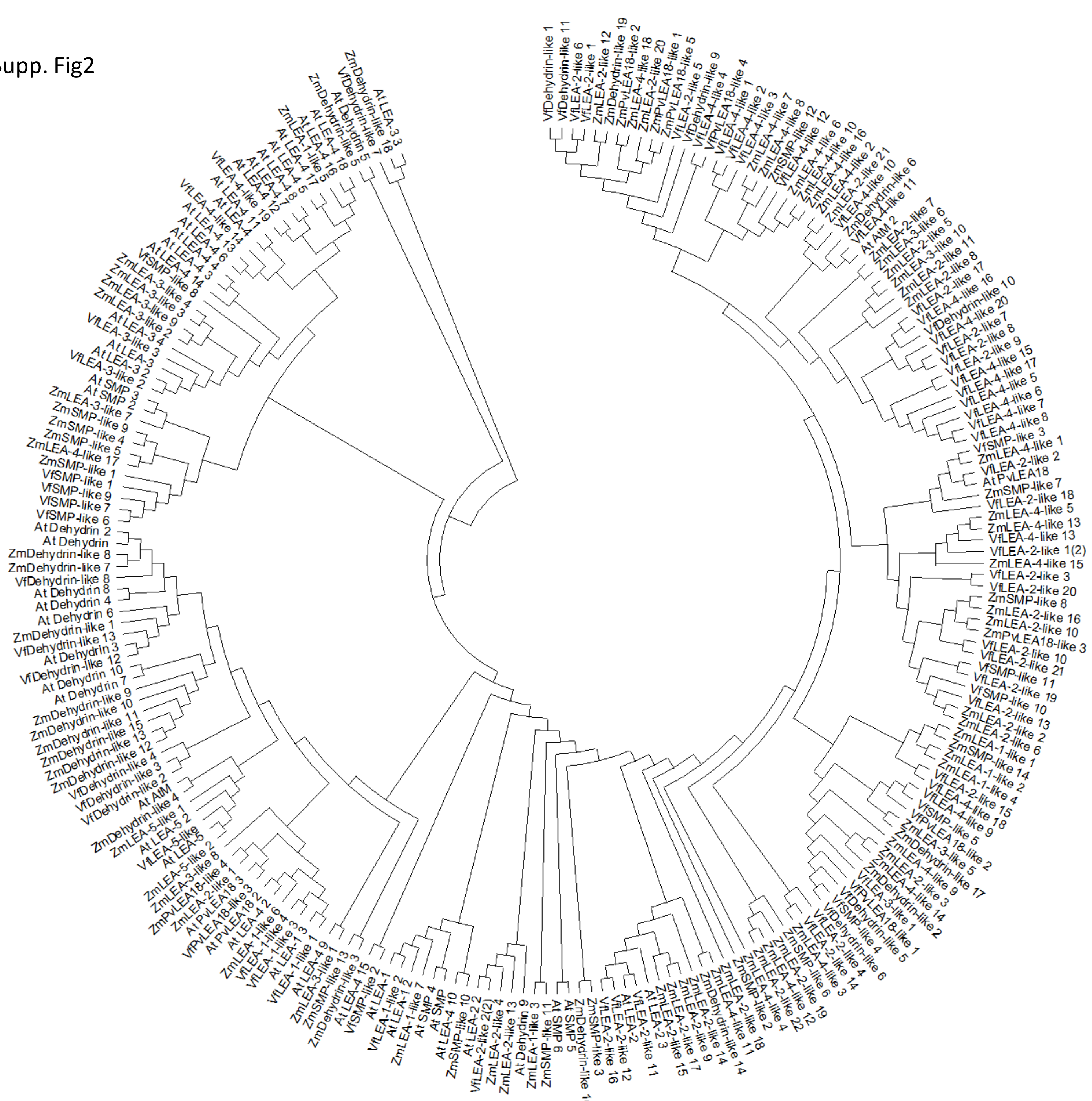

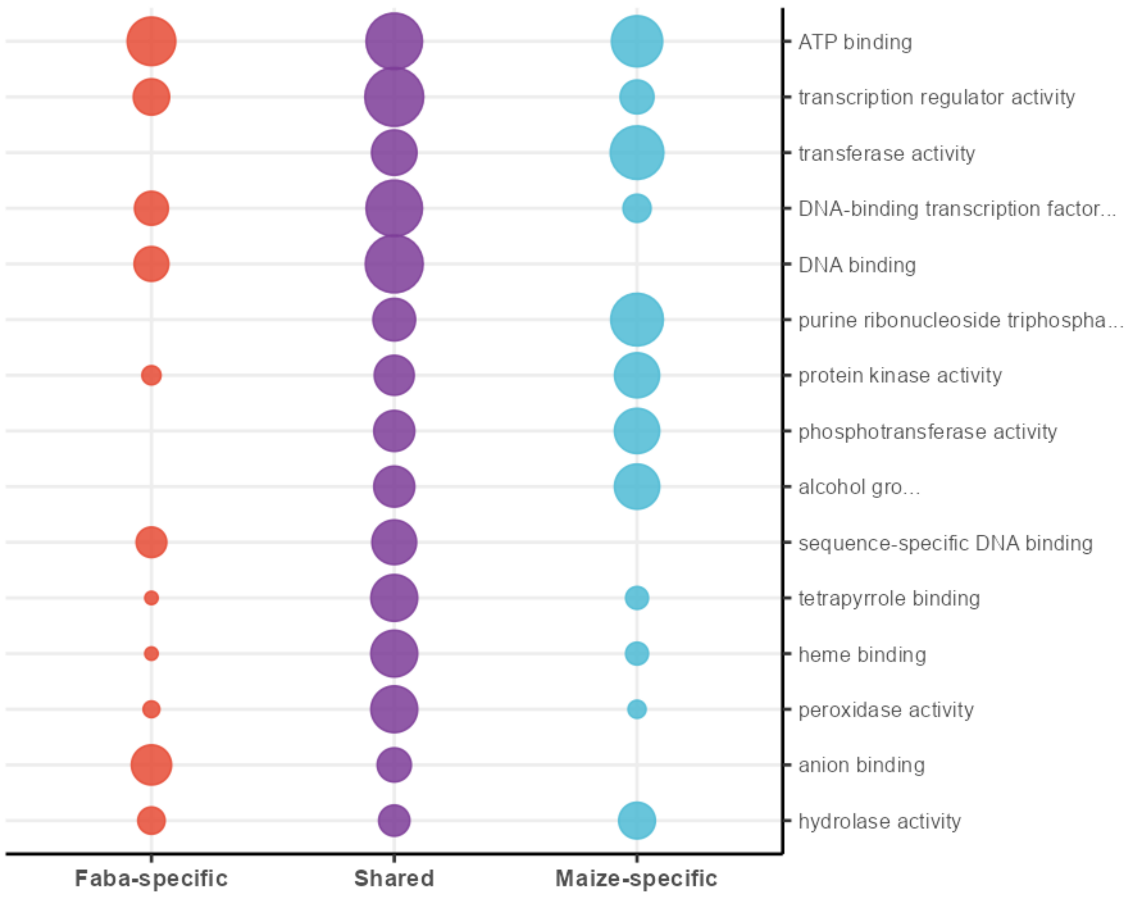
